# Osmolality controls the expression of cathelicidin antimicrobial peptide in human macrophages

**DOI:** 10.1101/332635

**Authors:** Youxian Li, Ingvild B. Johnsen

## Abstract

An imbalance between extracellular and intracellular fluid osmolality causes osmotic stress and affects cellular homeostasis. Recent research suggests that osmotic stress is also associated with various innate and adaptive immune responses. Here we present the surprising finding that osmolality tightly controls the expression of cathelicidin antimicrobial peptide (CAMP) in human macrophages. CAMP expression is strongly upregulated under hyperosmotic conditions and downregulated under hypoosmotic conditions. We also provide evidence that this osmolality-mediated antimicrobial response is dependent on nuclear factor of activated T-cells 5 (NFAT5) and mitogen-activated protein kinase (MAPK) p38. Finally, Toll-like receptor (TLR) activation inhibits osmolality-mediated expression of CAMP in human macrophages, suggesting that this osmolality-dependent regulation of CAMP is more relevant under homeostatic conditions, rather than during acute infections. This study expands our knowledge of the regulation of human antimicrobial peptides and highlights osmolality as an important and independent factor shaping host innate immune homeostasis.

## 1 Introduction

Osmolality describes the number of solute molecules per solution weight. In biological systems, the semipermeable membranes separate intracellular and extracellular fluid with distinct compositions. Hyperosmotic stress can have detrimental effects on cell homeostasis, causing water efflux and cell shrinkage, oxidative stress, DNA damage, cell cycle delay and cell deaths (Burg, Ferraris et al. 2007, Kuper, Beck et al. 2007, Brocker, Thompson et al. 2012). Mammalian cells have developed various osmoadaptive responses to compensate for the adverse effects due to solute concentration asymmetry. For example, under hyperosmotic conditions the transcription factor nuclear factor of activated T cells-5 (NFAT5) is activated to induce the expression of genes responsible for the synthesis or transport of uncharged small organic osmolytes such as sorbitol, betaine, and myo-inositol. These organic osmolytes can be accumulated to a high level to equalize intracellular and extracellular osmolality without perturbing macromolecular structure and function (Burg, Kwon et al. 1997).

The issue of hyperosmotic stress responses is well characterized in renal inner medulla, where cells are routinely exposed to high and varied levels of sodium and urea due to the operation of urinary concentrating mechanisms (Burg, Kwon et al. 1997, López-Rodríguez, Antos et al. 2004, Burg, Ferraris et al. 2007). However, recent studies provide evidence that osmolality is associated with various immune responses. It has been demonstrated that high salt promotes the induction of pathogenic Th17 cells (Kleinewietfeld, Manzel et al. 2013, Wu, Yosef et al. 2013), and dampens the immune suppressive functions of Treg cells (Hernandez, Kitz et al. 2015). In macrophages, hyperosmotic stress (or high sodium in particular) has been shown to promote the expression of pro-inflammatory genes and the production of nitric oxide (NO) upon LPS stimulation (Jantsch, Schatz et al. 2015, Zhang, Zheng et al. 2015, Tubbs, Liu et al. 2017), activate NLRP3 and NLRC4 inflammasomes (Ip and Medzhitov 2015), enhance type I interferon signaling (Zhang, Du et al. 2018) and reduce IL-4-and IL-13-mediated alternative (M2) activation (Binger, Gebhardt et al. 2015).

Antimicrobial peptides (AMPs) such as defensins and cathelicidins comprise a major component of innate immunity in mammals (Hilchie, Wuerth et al. 2013, Hancock, Haney et al. 2016). Human cathelicidin antimicrobial peptide (CAMP), also known as hCAP-18/LL-37/FALL-39, is the only known human cathelicidin. Human CAMP gene encodes the full-length 18k Da precursor, hCAP-18, which contains a conserved cathelin domain at the N-terminal and the functional antimicrobial domain at the C-terminal (Vandamme, Landuyt et al. 2012). The mature form of CAMP (LL-37) is released from hCAP-18 precursor through a proteinase 3-mediated cleavage process (Sørensen, Follin et al. 2001). CAMP has been shown to possess direct antimicrobial activities as well as various immunomodulatory properties (Yang, Chen et al. 2000, Scott, Davidson et al. 2002, Koczulla, Von Degenfeld et al. 2003, Vandamme, Landuyt et al. 2012). In humans CAMP expression is directly regulated by vitamin D signaling (Wang, Nestel et al. 2004, Gombart, Borregaard et al. 2005), although vitamin D-independent regulatory mechanisms have also been proposed (Park, Elias et al. 2011, Guo, Rosoha et al. 2013).

In this short report, we show that CAMP expression is tightly regulated by osmolality in human macrophages and reveal that NFAT5 and p38 kinase are the key components of the signaling pathway. We also demonstrate that canonical Toll-like receptor (TLR) activation interferes with this osmolality-driven antimicrobial response. Our study opens up a new understanding of innate immune signaling, demonstrating for the first time that a change in environmental osmolality alone is sufficient to elicit an antimicrobial response in human macrophages.

## Results

### Osmolality controls CAMP expression in human macrophages

We cultured human monocyte-derived macrophages (MDMs) under hyperosmotic (adjusted by NaCl or sucrose) or hypoosmotic conditions (adjusted by H_2_O) to evaluate the effect of osmolality on CAMP expression. Active vitamin D (1,25(OH)_2_D_3_), a known inducer of CAMP expression in human macrophages (Liu, Stenger et al. 2006, Yuk, Shin et al. 2009), was used as the positive control. To avoid any effect mediated by serum vitamin D all cells were cultured under serum-free conditions in this study. As shown in Figure 1A-C, hyperosmolality upregulated CAMP mRNA and protein expression in a dose-dependent manner. Notably, a modest increase in osmolality by 50mOsm or 100mOsm was sufficient to induce higher levels of CAMP comparing to 1,25(OH)_2_D_3_ treatment. NaCl and sucrose showed similar inductive effects, indicating that this osmolality-mediated CAMP response is independent of the types of osmolytes. Interestingly, CAMP expression was reduced in macrophages with hypoosmotic treatment, suggesting that osmolality contributes to basal CAMP expression even at physiologically isosmotic conditions. Taken together these data indicate that osmolality effectively controls CAMP expression in human macrophages.

**Figure 1:**
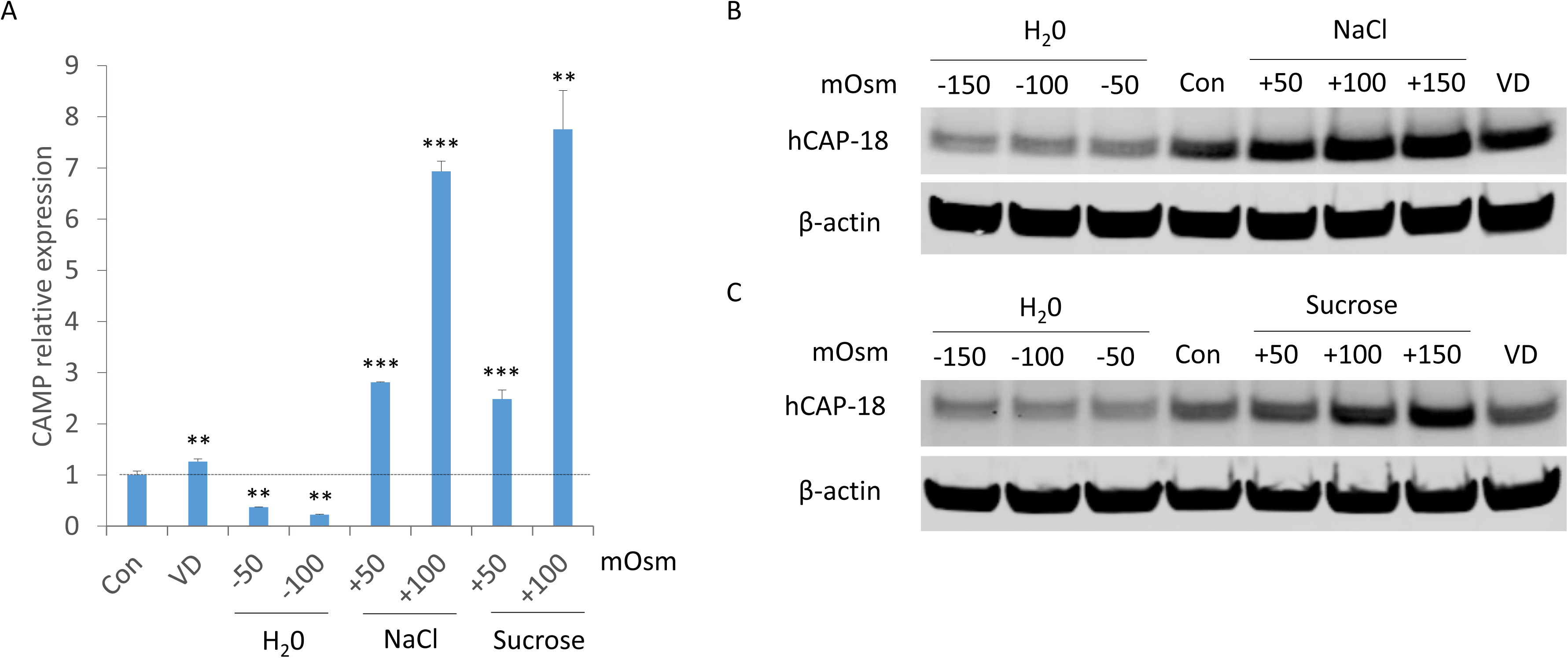
Osmolality controls CAMP expression in human macrophages. (A-C) MDMs were cultured in control medium with no osmolality adjustment (Con), hyperosmotic medium (adjusted by NaCl or sucrose), hypoosmotic medium (adjusted by H_2_O), or treated with 100nM 1,25(OH)_2_D_3_ (VD) for 24 hours. CAMP mRNA (A) and protein expression (hCAP-18) (B, C) was assessed by qRT-PCR and Western Blot. Error bars represent SD for triplicates. ^**^p < 0.01, ^***^p < 0.001 (compared to Con). Data representative of at least three independent experiments from different donors.

### Osmolality-mediated CAMP expression in human macrophages are regulated by NFAT5 and p38

We went on to investigate the molecular mechanisms underlying osmolality-mediated CAMP response. As NFAT5 is a central component of osmoadaptive signaling, we used NFAT5 siRNA (siNFAT5) to knock down NFAT5 gene expression before we manipulated medium osmolality. siNFAT5 effectively suppressed NFAT5 gene expression before hyperosmotic treatment (Figure 2A). Prior knockdown of NFAT5 expression strongly repressed basal and hyperosmolality-induced CAMP expression (Figure 2B, C), indicating that CAMP expression is controlled by osmolality via a NFAT5-dependent mechanism. Interestingly, hyperosmotic treatment did not enhance NFAT5 expression in control siRNA-treated cells, but promoted NFAT5 expression in siNFAT5-treated cells (Supplementary Figure 1A, B). This suggests that the constitutive expression of NFAT5 in human macrophages is normally sufficient, but a feedback loop exists so that when cells with insufficient NFAT5 levels sense the need to elicit osmoadaptive responses, NFAT5 expression can be levelled up.

**Figure 2:**
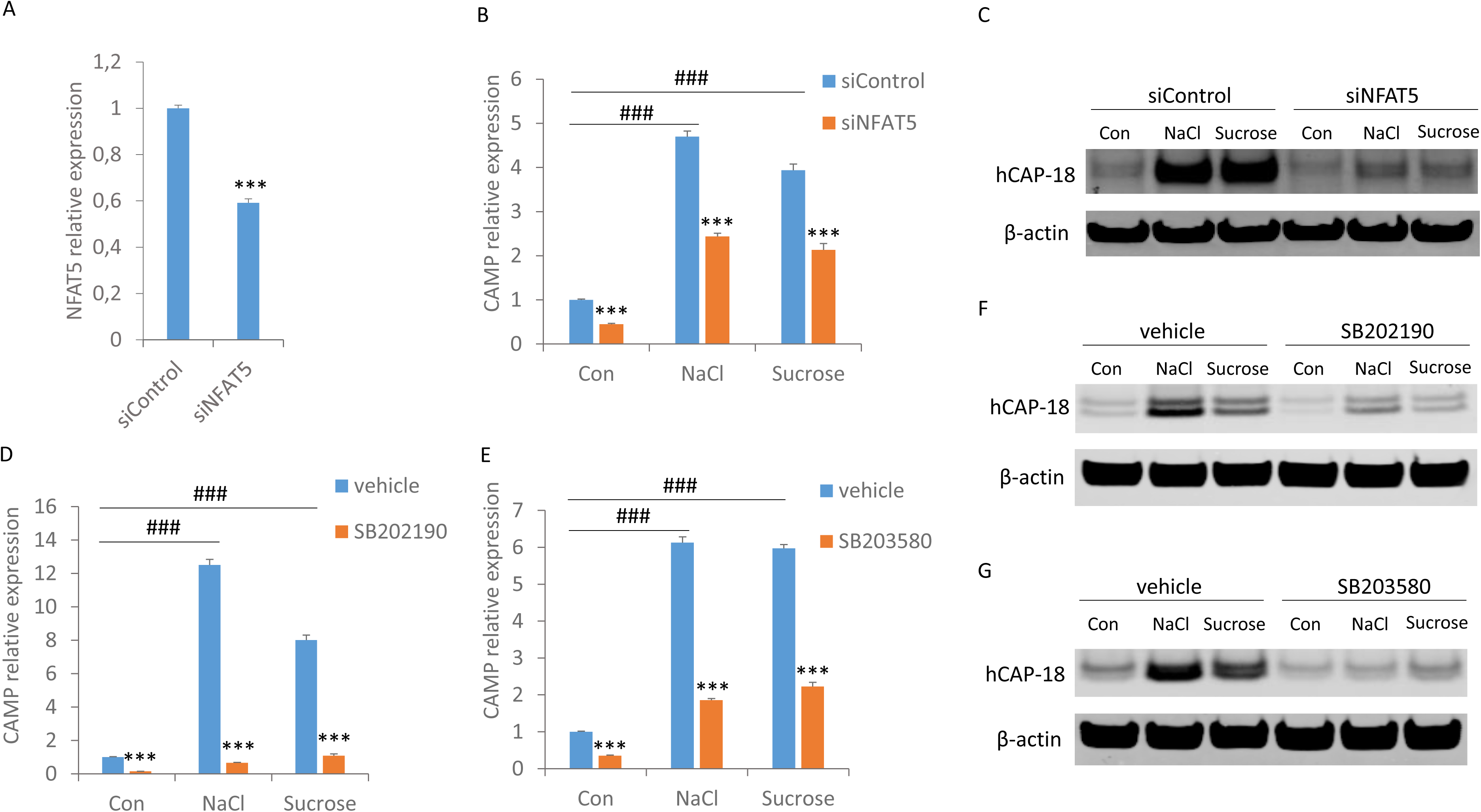
Osmolality-mediated CAMP expression in human macrophages are regulated by NFAT5 and p38. (A) MDMs were transfected with siRNA targeting NFAT5 (siNFAT5) or scrambled control siRNA (siControl) for 72 hours. NFAT5 mRNA expression was assessed by qRT-PCR. (B, C) MDMs were transfected with siRNA targeting NFAT5 (siNFAT5) or scrambled control siRNA (siControl) for 72 hours before cultured in control medium with no osmolality adjustment (Con), or medium with additional 100mOsm NaCl or sucrose for 24h. CAMP mRNA expression (B) was assessed by qRT-PCR and CAMP (hCAP-18) protein expression was assessed by Western Blot (C). (D-G) MDMs were pretreated with vehicle, or p38 kinase inhibitor SB202190 (D, F) or SB203580 (E, G) for 30min, then cultured in control medium with no osmolality adjustment (Con), or medium with additional 100mOsm NaCl or sucrose. CAMP mRNA (D, E) and protein expression (hCAP-18) (F, G) was assessed by qRT-PCR and Western Blot. Error bars represent SD for triplicates. ^***^p < 0.001 (comparing to corresponding siControl or vehicle samples). ###p < 0.001. Data are representative of at least three independent experiments from different donors.

The molecular mechanisms underlying NFAT5 activation is not well understood, however it has been shown that p38, a subgroup of mitogen-activated protein kinases (MAPKs), is required for osmolality-dependent activation of NFAT5 (Ko, Lam et al. 2002, Zhou 2015, Zhou 2016). Therefore, we examined if p38 contributes to osmolality-meidated CAMP expression. We used two selective p38 kinase inhibitors, SB202190 and SB203580 (Wilson, McCaffrey et al. 1997, Young, McLaughlin et al. 1997), to block p38 kinase activity. SB202190 and SB203580 pretreatment strongly reduced basal and hyperosmolality-induced CAMP expression both at the mRNA level (Figure 2D, E) and at the protein level (Figure 2F, G), indicating that p38 kinase activity is also important for the signaling of osmolality-mediated CAMP expression.

### Toll-like receptor (TLR) activation inhibits osmolality-mediated CAMP expression in human macrophages

High salt has been reported to potentiate the expression of various cytokines and pro-inflammatory genes when macrophages are activated by TLR ligands such as LPS (Jantsch, Schatz et al. 2015, Zhang, Zheng et al. 2015, Tubbs, Liu et al. 2017). We next asked if TLR stimulation affects osmolality-mediated CAMP response in human macrophages. We treated macrophages with two TLR ligands: Pam3CSK4 (a TLR1/2 ligand) and LPS (a TLR4 ligand), and cultured the cells under isosmotic or hyperosmotic conditions. Interestingly, both Pam3CSK4 and LPS treatment strongly reduced basal and hyperosmolality-induced CAMP expression (Figure 3A-C). Pam3CSK4 and LPS did not downregulate NFAT5 gene expression in macrophages (Figure 3D), suggesting that the suppressive effect of TLR agonists may be modulated via post-transcriptional regulation of NFAT5 (e.g., inhibition of NFAT5 protein activation), or via a NFAT5-independent mechanism. An earlier report (Dhawan, Wei et al. 2015) and our recent study (Li, Østerhus et al. 2018) have shown that the transcription factor C/EBPα plays a critical role in mediating CAMP expression in human airway epithelial cells and macrophages. Therefore, we examined the expression of C/EBPα in macrophages upon Pam3CSK4 or LPS treatment. As shown in Figure 3E, C/EBPα expression was repressed by LPS or Pam3CSK4 stimulation, corresponding to downregulated CAMP expression (Figure 3A-C). Taken together our data suggest that TLR activation suppresses osmolality-mediated CAMP expression possibly via downregulation of C/EBPα.

**Figure 3:**
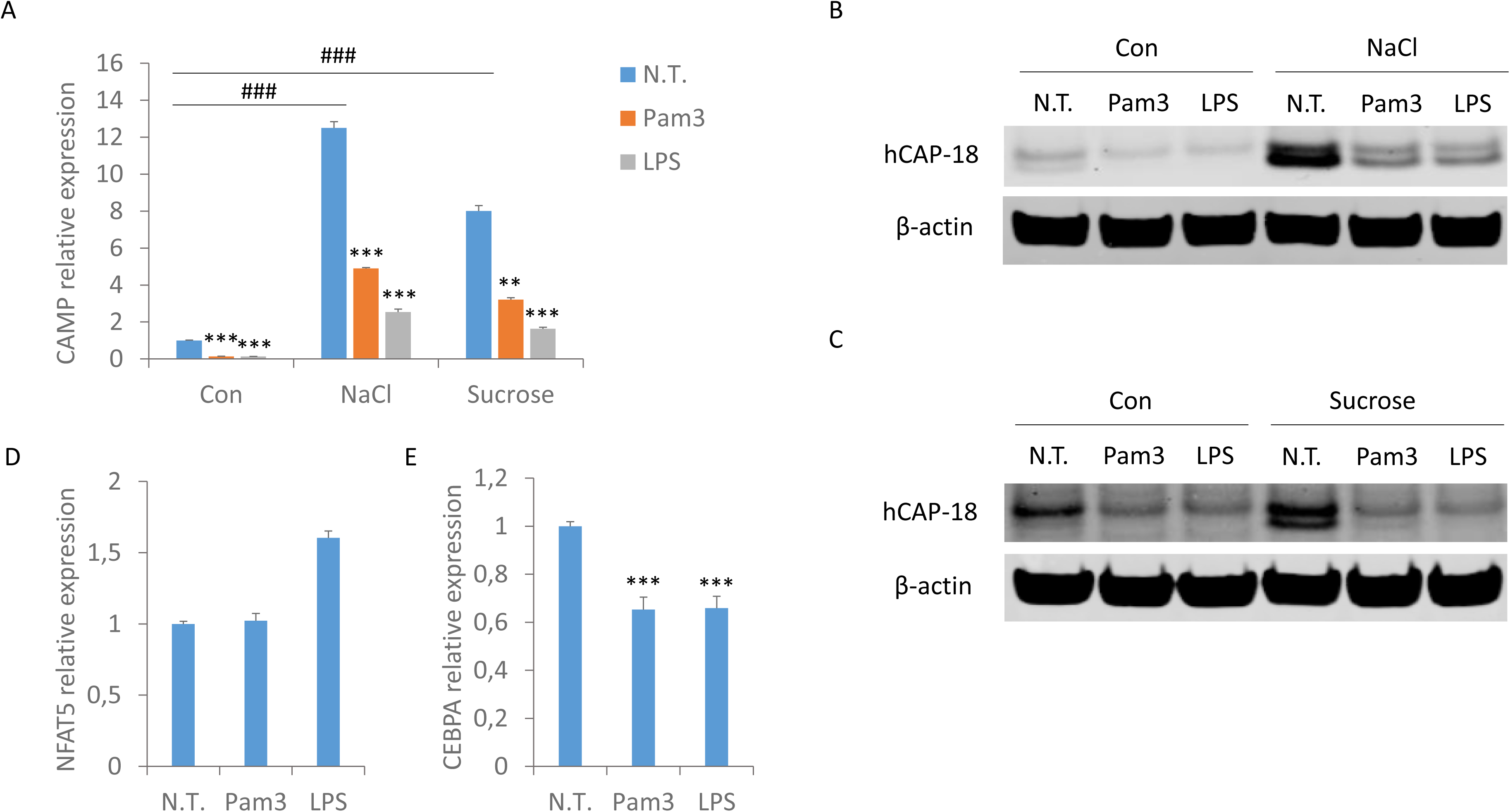
Toll-like receptor (TLR) activation inhibits osmolality-mediated CAMP expression in human macrophages. (A-C) MDMs were cultured in control medium with no osmolality adjustment (Con), or medium with additional 100mOsm NaCl or sucrose, concomitant with mock treatment (N.T.) or treatment with 1µg/mL Pam3CSK4 (Pam3) or LPS. CAMP mRNA (A) and protein expression (hCAP-18) (B, C) was assessed by qRT-PCR and Western Blot. (D, E) MDMs were mock-treated (N.T.) or treated with 1µg/mL Pam3CSK4 (Pam3) or LPS, NFAT5 (D) and C/EBPα (E) mRNA expression was assessed by qRT-PCR. Error bars represent SD for triplicates. ^**^p < 0.01, ^***^p < 0.001 (comparing to corresponding N.T. samples); ###p < 0.001. Data are representative of at least three independent experiments from different donors.

## Discussion

The molecular mechanisms underlying macrophage activation and defense gene expression have been extensively studied over the last two decades. Although the canonical structural recognition of pathogen-associated molecular patterns (PAMPs) or danger-associated molecular patterns (DAMPs) and the downstream signaling pathways are well-characterized, accumulating evidence suggests that macrophages can also sense and respond to environmental changes, such as nutrient availability and oxygen levels (Strehl, Fangradt et al. 2014, O’Neill and Pearce 2016, Langston, Shibata et al. 2017). The association between osmolality and macrophage functions is only beginning to be appreciated (Binger, Gebhardt et al. 2015, Ip and Medzhitov 2015, Jantsch, Schatz et al. 2015, Zhang, Zheng et al. 2015, Tubbs, Liu et al. 2017, Zhang, Du et al. 2018). These studies suggest that hyperosmotic stress (or high NaCl specifically) tends to polarize macrophages towards a more pro-inflammatory phenotype. Our report however indicates that osmolality is more than a factor potentiating inflammation. We provide clear evidence that human macrophages can effectively monitor osmolality changes in the environment to elicit (or repress) an important antimicrobial response, in the absence of other inflammatory stimuli.

The following observations make this newly identified regulatory mechanism of CAMP particularly intriguing. First, osmolality can both positively and negatively affect CAMP expression: hyperosmolality upregulates CAMP expression whilst hypoosmolality downregulates CAMP expression, suggesting that this is a homeostatic regulatory mechanism adapting macrophage CAMP expression levels to osmolality changes in the environment. Secondly, CAMP expression is very sensitive to changes in osmolality: a modest increase by 50 or 100mOsm induces higher expression of CAMP comparing to active vitamin D treatment. Thirdly, hyperosmolality induces CAMP expression via a shared mechanism regardless of the types of osmolytes, as both the ionic and non-ionic osmolytes (NaCl and sucrose) potently induce CAMP expression via the same NFAT5-and p38-dependent signaling pathway. Finally, hyperosmolality alone is sufficient to drive CAMP expression without the need of additional stimulation (e.g., LPS treatment). In fact CAMP expression is strongly reduced when macrophages are primed by Pam3CSK4 or LPS, emphasizing that this osmolality-mediated CAMP response is a kind of homeostatic regulation (and possibly a preventive defense mechanism), rather than an acute response to infections. This observation supports what we have proposed earlier (Li, Østerhus et al. 2018), that human macrophages express CAMP under homeostatic conditions, but shift away from CAMP production to prioritize production of other effector molecules such as cytokines and chemokines when encountering a serious pathogenic threat.

The physiological implications of this osmolality-dependent regulation of CAMP remain to be elucidated. It will also be interesting to investigate if this regulatory mechanism of AMPs is present in species other than humans. Given the prominent antimicrobial property of CAMP, it is tempting to speculate that this regulatory mechanism may be particularly relevant for local defense in tissues with significant fluctuations in osmolality, such as the gut and the urinary system. Our study will likely provide useful insights into the maintenance and breakdown of immune homeostasis in these tissues. In addition, this newly identified osmolality-dependent regulation offers a potentially simple and practical approach to manipulate CAMP levels *in vivo*, either for investigative purpose to reveal CAMP functions that are truly physiologically relevant, or for potential therapeutic purposes to boost natural and homeostatic antimicrobial responses.

## Materials and methods Reagents

Active vitamin D (1,25(OH)_2_D_3_) was purchased from Tocris Bioscience and used at a working concentration of 100nM. Pam3CSK4 was purchased from Invivogen and used at a working concentration of 1µg/mL. LPS was purchased from Sigma Aldrich and used at a working concentration of 1µg/mL. SB202190 and SB203580 were purchased from Sigma Aldrich and used at working concentrations of 10 µM and 20 µM respectively (30 min pretreatment).

### Cell culture and medium osmolality adjustment

Peripheral blood mononuclear cells (PBMCs) were isolated from fresh buffy coats of healthy donors using gradient centrifugation with Lymphoprep^™^ (Axis-Shield). Buffy coats were supplied by the blood bank at St.Olavs Hospital in Trondheim, Norway, and their use in research has been approved by the Regional Committee for Medical and Health Research Ethics (REK), and by the donors themselves. Cells were washed with PBS and seeded in RPMI 1640 medium (supplemented with 0.34 mM L-glutamine and 10 µg/mL gentamicin). After two hours non-adherent cells were removed by washing with RPMI 1640 medium. Monocytes were cultivated in RPMI 1640 medium supplemented with 10% human serum (heat inactivated, obtained from blood bank of St. Olavs Hospital, Trondheim), 0.34 mM L-glutamine, 10 µg/mL gentamicin and 10 ng/mL M-CSF (Biolegend) for macrophage differentiation. Medium was changed every three days. Macrophages differentiated for 12-16 days were used in this study. On the day of treatment, medium was switched to serum-free RPMI 1640 (with or without osmolality adjustment). Medium osmolality was adjusted by sterilized deionized water, 300mM sucrose (in RPMI) or 150 mM NaCl (in RPMI). After 24h samples were collected for quantitative real-time PCR or Western blot analysis.

### Quantitative real-time PCR (qRT-PCR)

RNA was isolated with the RNeasy mini kit (Qiagen) following the manufacturer’s protocol. cDNA was synthesized from isolated RNA using the qScript kit (Quanta) following the manufacturer’s protocol. Quantitative real-time PCR (qRT-PCR) was performed using Perfecta SYBR Green reaction mix (Quanta) and a StepOnePlus instrument (Life Technologies) with the temperature profile 95°C for 20 s, 40 cycles at 95°C for 3s and 60°C for 30s. Fold change in gene expression was calculated using the ΔΔCt-method normalized against GAPDH. Sequences of primers used in this study are as follows: CAMP: forward 5′-TCGGATGCTAACCTCTACCG-3′, reverse 5′-GTCTGGGTCCCCATCCAT-3′; GAPDH: forward 5′-GAAGGTGAAGGTCGGAGTC-3′, reverse 5′-GAAGATGGTGATGGGATTTC-3′; CEBPA: forward 5′-GGAGCTGAGATCCCGACA-3′, reverse 5′-TTCTAAGGACAGGCGTGGAG-3′

### Western Blot

Cells were washed once in PBS and lysed in lysis buffer (50mM Tris-HCl, 150mM NaCl, 10% Glycerol, 0,5% Triton X-100 and 2mM EDTA) containing phosphatase and protease inhibitors (100 mM Sodium Fluoride, 1 mM Sodium Orthovanadate, 40 mM β-Glycerophosphate, 10 µg/mL Leupeptin, 1 µM Pepstatin A and 1 mM Phenylmethylsulfonyl fluoride). Protein extracts were separated by NuPAGE^®^ Bis-Tris gels (Thermo Fisher Scientific) and dry blotting was performed using iBlot^®^ Gel Transfer stacks Nitrocellulose Mini kit and iBlot^®^ machine (Invitrogen). Primary human antibodies for hCAP-18 (#650302) and NFAT5 (#F-9) were purchased from Biolegend and Santa Cruz respectively. Household β-actin antibody (A1978) was purchased from Sigma Aldrich and used as a loading control. Secondary antibodies (IRDye^®^ 800CW Goat anti-Mouse, IRDye^®^ 680RD Goat anti-Mouse) were purchased from LI-COR Biosciences. LICOR Odyssey imager was used as the scanning system.

### RNA interference

siRNAs were purchased from Qiagen (AllStars control siRNA) and Ambion (NFAT5) respectively. siRNAs (20nM) were reverse transfected into cells using Lipofectamine RNAiMAX (Thermo Fisher Scientific) transfection reagent according to the manufacturer’s instructions. Cells were incubated with siRNA-RNAiMAX complex for 72 hours before further treatment.

### Statistics

Results are expressed as mean + SD (n=3). A two-sided P-value < 0.05 as determined by Student t-test was considered significant. All data are representative of at least three independent experiments with PBMCs from different donors.

## Data availability

Data supporting the conclusions of this manuscript will be made available by the authors, without undue reservation, to any qualified researcher.

## Acknowledgements

The work was funded by the Research Council of Norway (grant number 230381). We thank Kristin Rian for excellent technical support.

## Author contributions statement

Y.L. conducted the experiments. Y.L. and I.B.J. analyzed the results. Y.L. wrote the manuscript. Both authors reviewed and approved the manuscript.

## Ethics Statement

Buffy coats used in this study were supplied by the blood bank at St. Olavs Hospital in Trondheim, Norway, and their use in research has been approved by the Regional Committee for Medical and Health Research Ethics (REK), and by the donors themselves.

**Figure S1:**
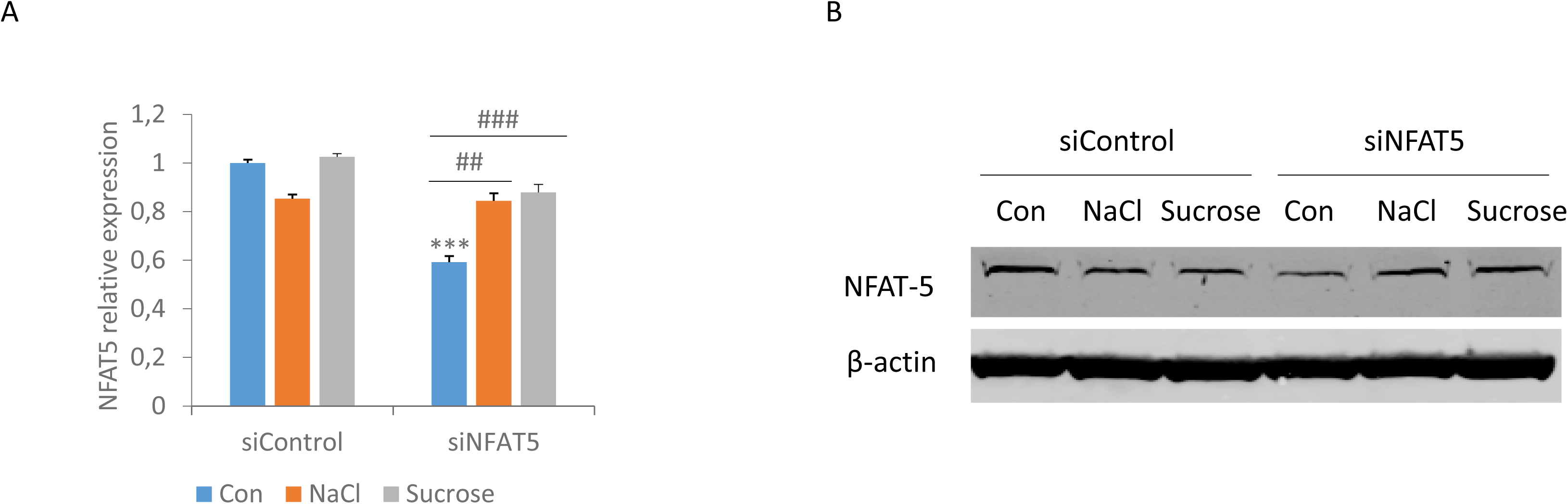
Hyperosmolality does not enhance constitutive NFAT5 expression, but reverses the suppressive effect of NFAT5 siRNA in human macrophages. (A, B) MDMs were transfected with siRNA targeting NFAT5 (siNFAT5) or scrambled control siRNA (siControl) for 72 hours before cultured in control medium with no osmolality adjustment (Con), or medium with additional 100mOsm NaCl or sucrose for 24h. NFAT5 mRNA expression (A) was assessed by qRT-PCR and CAMP (hCAP-18) protein expression was assessed by Western Blot (B). ^***^p < 0.001 (compared to the corresponding siControl sample). ##p < 0.01. ###p < 0.001. Data representative of at least three independent experiments from different donors.

